# Engineered Cpf1 Enzymes with Altered PAM Specificities

**DOI:** 10.1101/091611

**Authors:** Linyi Gao, David B.T. Cox, Winston X. Yan, John Manteiga, Martin Schneider, Takashi Yamano, Hiroshi Nishimasu, Osamu Nureki, Feng Zhang

## Abstract

The RNA-guided endonuclease Cpf1 is a promising tool for genome editing in eukaryotic cells^1-5^. Compared to other genome editing platforms, Cpf1 offers distinct advantages, such as the ability to easily target multiple genes simultaneously^3^, as well as low rates of off-target activity^4, 5^. However, the *Acidaminococcus sp. BV3L6* Cpf1 (AsCpf1), which has been successfully harnessed for genome editing, can only robustly cleave target sites preceded by a TTTV protospacer adjacent motif (PAM), which may limit its practical utility. To address this limitation, we used a structure-guided saturation mutagenesis screen to increase the targeting range of Cpf1. We engineered two variants of AsCpf1 with the mutations S542R/K607R and S542R/K548V/N552R that can cleave target sites with TYCV/CCCC and TATV PAMs, respectively, with enhanced activities *in vitro* and in human cells. Genome-wide assessment of off-target activity indicated that these variants retain a high level of DNA targeting specificity, which can be further improved by introducing mutations in non-PAM-interacting domains. Together, these variants increase the targeting range of AsCpf1 to one cleavage site for every ~8.7 bp in non-repetitive regions of the human genome, providing a useful addition to the CRISPR/Cas genome engineering toolbox.

---

The RNA-guided endonuclease Cpf1 from microbial CRISPR-Cas adaptive immune systems is a promising tool for genome editing in eukaryotic cells^1^. Compared to other genome editing platforms such as ZFNs, TALENs, and CRISPR-Cas9, Cpf1 offers distinct advantages. Cpf1 is naturally a single RNA-guided endonuclease and can process its own crRNA array into mature crRNAs to facilitate targeting of multiple genes simultaneously^2, 3^. Moreover, Cpf1 has low mismatch tolerance and high levels of on-target specificity^4, 5^.

Previously we have identified two orthologs of Cpf1 with robust activity in mammalian cells, *Acidaminococcus sp. BV3L6* Cpf1 (AsCpf1) and *Lachnospiraceae bacterium ND2006* Cpf1 (LbCpf1)^1^, both of which require a TTTV protospacer adjacent motif (PAM), where V = A, C, or G. This motif corresponds, on average, to one cleavage site for every 23 base pairs (bp) in non-repetitive regions of the human genome. For applications for which the location of the target site is critical, *e.g.* homology-directed repair or loss-of-function mutation generation at specific exonic locations, the requirement of a TTTV PAM may limit the availability of suitable target sites, reducing the practical utility of Cpf1. To address this limitation, we aimed to engineer variants of Cpf1 that can recognize alternative PAM sequences in order to increase its targeting range.

Previous work has shown that the PAM preference of Cas9 can be altered by mutations to residues in close proximity with the PAM DNA duplex^6-9^. We sought to investigate whether the PAM preference of Cpf1, despite its strong evolutionary conservation^1^, can also be modified. Based on the crystal structure of AsCpf1 in complex with crRNA and target DNA^10^, we selected 64 residues in AsCpf1 in proximity to the PAM duplex (Fig. 1a) for targeted mutagenesis. By randomizing the codons at each position using cassette mutagenesis11, we constructed a plasmid library of AsCpf1 variants encoding all possible single amino acid substitutions at these residues. To assay cleavage activity, we adapted a plasmid interference-based depletion screen in *E. coli*^1, 6, 12, 13^ to identify variants within this library with cleavage activity at non-canonical PAMs (Fig. 1b). In our modified assay, a pool of *E. coli*, each expressing crRNA and a variant of Cpf1, was transformed with a plasmid carrying an ampicillin resistance gene and a target site bearing a mutated PAM. Cleavage of the target resulted in the loss of ampicillin resistance and subsequent cell death when grown on ampicillin selective media. By sequencing the plasmid DNA in surviving bacteria, we identified the variants that were depleted; these variants were active at the mutant PAM.

**Figure 1.**
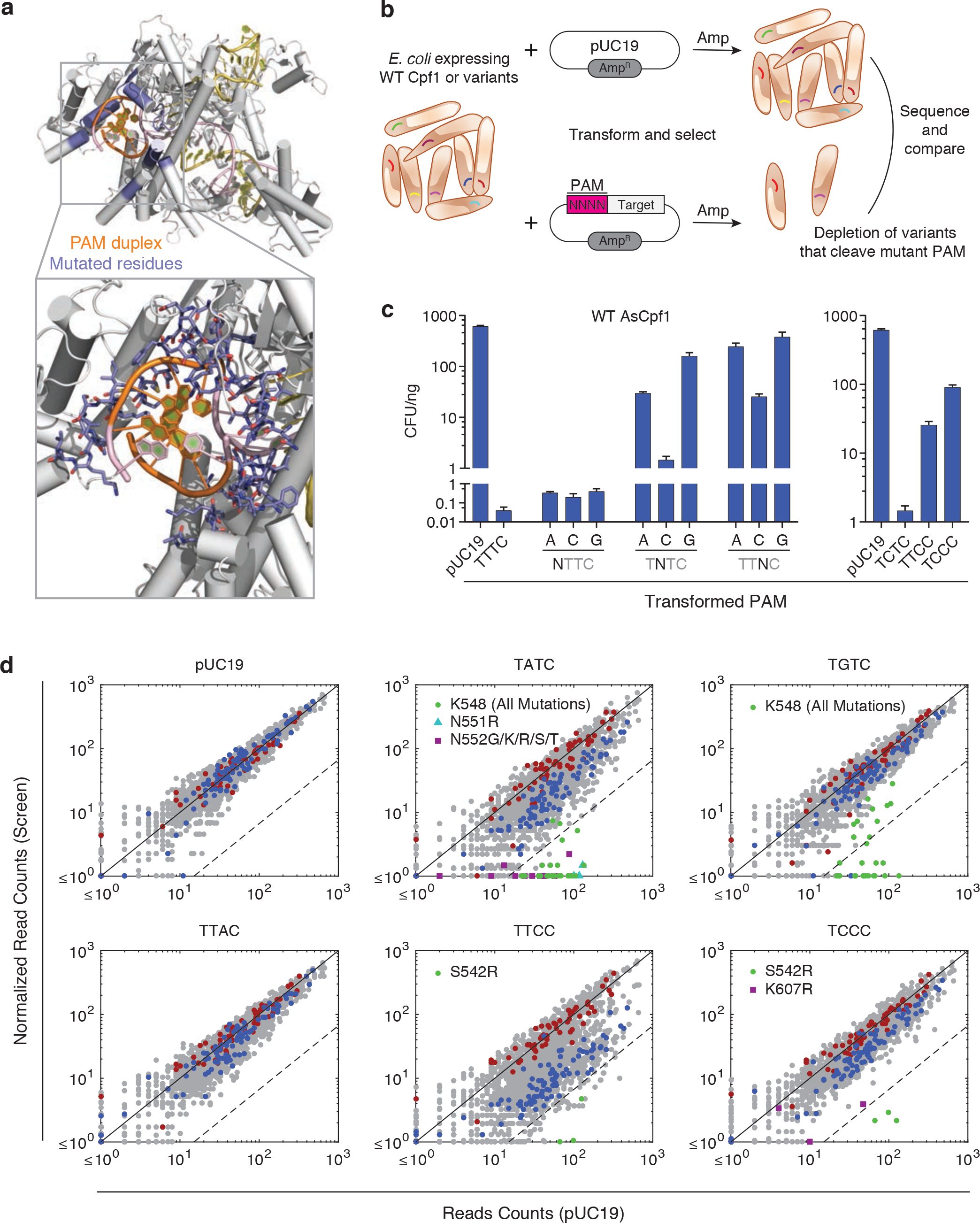
A bacterial interference-based negative selection screen identifies amino acid substitutions of Cpf1 conferring activity at non-canonical PAMs. (**a**) Crystal structure of AsCpf1 in complex with crRNA and target DNA, showing the PAM duplex (orange), and mutated residues (blue). (**b**) Schematic of bacterial interference assay used for depletion screening for Cpf1 variants with altered PAM specificity. (**c**) Sensitivity of wild-type AsCpf1 to substitutions mutations in the PAM. (**d**) Screen readout, highlighting depleted hits. Each dot represents a distinct Cpf1 wild-type (WT) or mutant codon. The dashed line indicates 15-fold depletion. Red = stop codon; blue = WT codon.

In order to use this assay to distinguish the variants from WT AsCpf1, we first evaluated the sensitivity of WT AsCpf1 to substitution mutations in the PAM, as determined by plasmid depletion in *E. coli* due to successful plasmid interference. We focused on PAMs with single nucleotide substitutions (*i.e.*, NTTV, TNTV, and TTNV, where V was arbitrarily chosen to be C). When transformed with NTTC and TCTC PAMs, *E. coli* expressing WT AsCpf1 had negligible survival (Fig. 1c), indicating that these PAM sequences supported AsCpf1-mediated DNA plasmid cleavage and were not usable for screening the variant library. In contrast, the other five PAMs with a single mutation (TATC, TGTC, TTAC, TTCC, and TTGC) had notable survival rates. We subsequently screened the variant library for activity at these five PAMs, as well as an additional PAM with a double mutation (TCCC) (Fig. 1d and **Supplementary Fig. 1a**).

Following sequencing readout, approximately 90% of the variants in the library were present in enough abundance in the negative control (minimum ~15 reads) to assess depletion. For TATC, TGTC, TTCC, and TCCC PAMs, at least one AsCpf1 variant in the library was highly depleted (≥15-fold) (Fig. 1d). For TATC and TGTC, the majority of depleted hits were amino acid substitutions at K548, a conserved residue that forms hydrogen bonds with the second base of the PAM^10, 14^. A smaller number of hits were observed for TTCC and TCCC, most notably an arginine substitution at S542, a non-conserved residue.

We evaluated whether hits identified in the screen had activity in HEK293FT cells (Fig. 2a and **2b**). Most of the hits tested generated indels for their corresponding PAMs; in particular, K548V was most active at a TATC genomic target site, while S542R dramatically increased activity for both a TTCC and a TCCC target site. Combining the top single amino acid mutations into double and triple mutants further improved activity (Fig. 2b and **Supplementary Fig. 1b**). We selected the variants with the highest activity, S542R/K607R (hereafter referred to as RR) and S542R/K548V/N552R (hereafter referred to as RVR), for further investigation.

**Figure 2.**
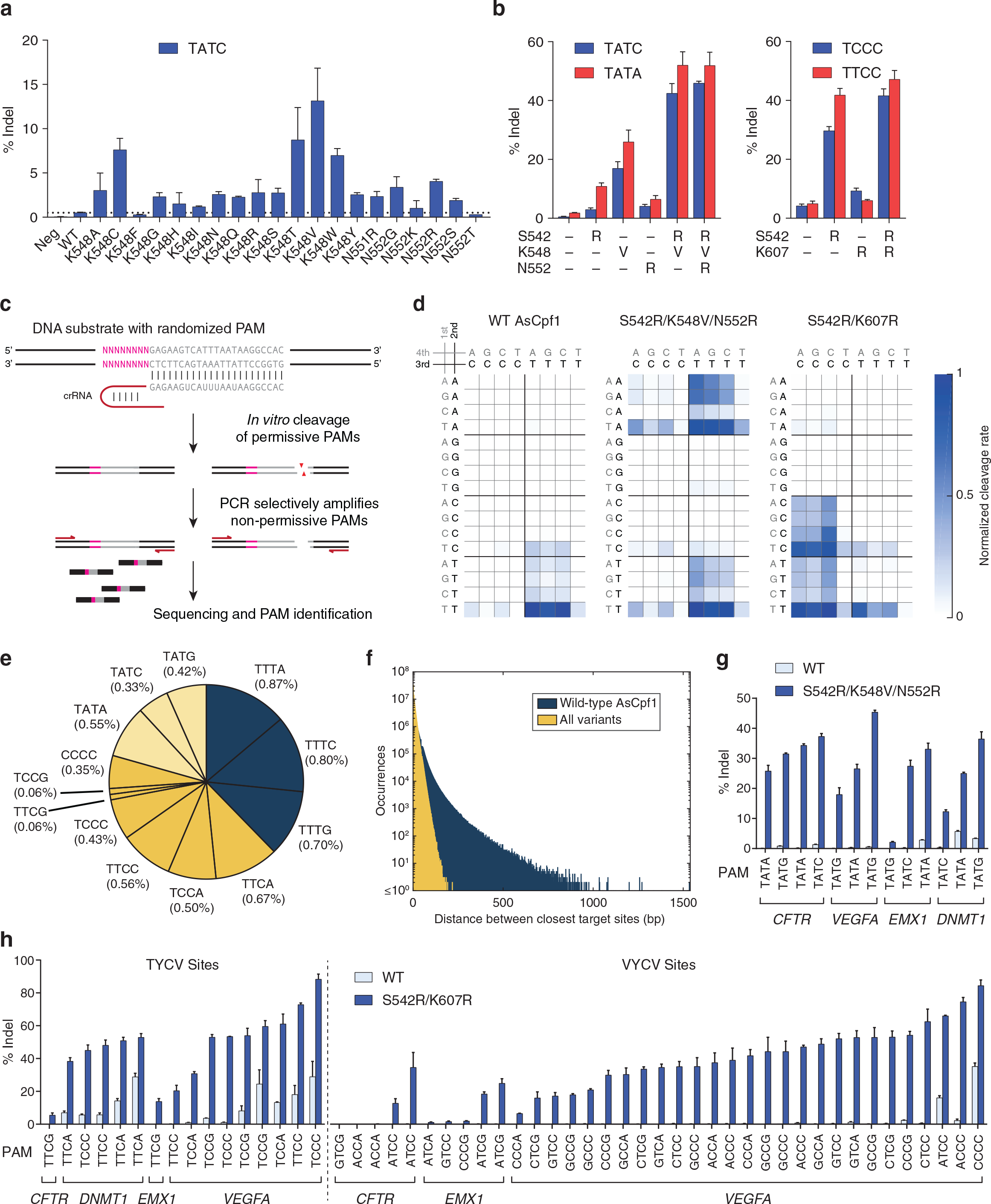
Construction and characterization of Cpf1 variants with altered PAM specificities. (**a**) Evaluation of single amino substitution hits for a target site with a TATC PAM. (**b**) Combinatorial mutagenesis to construct Cpf1 mutants with robust activity at target sites bearing TATV and TYCV PAMs. (**c**) Schematic of *in vitro* cleavage experiment used to determine PAM specificities. (**d**) Normalized cleavage rates for all 4-base PAM motifs for wild-type, S542R/K607R, and S542R/K548V/N552R variants. NNRN PAMs are not shown due to negligible cleavage. (**e**) Targeting range of Cpf1 variants in the human genome, including WT (dark blue), S542R/K607R (yellow), and S542R/K548V/N552R (light yellow). The percentages indicate the proportion of all non-repetitive guide sequences (both top and bottom strands) represented by the corresponding PAM. (**f**) Distance between nearest target sites in non-repetitive regions of the human genome for TTTV PAMs (dark blue) and all PAMs cleavable by any of the variants (yellow). (**g**) Activity of the S542R/K548V/N552R variant at TATV target sites. (**h**) Activity of the S542R/K607R variant at TYCV and VYCV target sites. All indel percentages were measured in HEK293 cells.

To assess the global PAM preference of the RR and RVR variants and to compare with the WT, we adapted an *in vitro* PAM identification assay described previously (Fig. 2c)^1, 15^. We incubated cell lysate from HEK293 cells expressing AsCpf1 (or a variant) with *in vitro*-transcribed crRNA and a library of plasmid DNA containing a constant target preceded by a degenerate sequence (5’- NNNNNNNN-target). For each Cpf1 variant, ten replicates of the cleavage reaction were carried out, each incubated for a different amount of time, in order to determine cleavage kinetics (see **Methods**). As expected, WT AsCpf1 was most active at TTTV PAMs (Fig. 2d), although we also observed cleavage at low rates for NTTV, TCTV, TTCV, and TTTT, consistent with our observations in HEK293 cells (**Supplementary Fig. 1c**). In contrast, the RR and RVR variants had the highest activity at TYCV/CCCC and TATV PAMs, respectively, compared to little or no activity for WT at those PAMs (Fig. 2d). Collectively, these new PAMs increase the targeting range of Cpf1 by 2.6-fold in non-repetitive regions of the human genome (Fig. 2e) and reduce the sizes of genomic DNA stretches that cannot be targeted (Fig. 2f).

Next, we investigated the activity of the RR and RVR variants at their preferred PAMs across a larger panel of target sites in HEK293 cells (Fig. 2g and **2h**). Both variants mediated robust editing across the four genes assessed (*CFTR*, *DNMT1*, *EMX1*, and *VEGFA*); in particular, 14/16 (88%) of TYCV target sites had >20% indel for RR (vs. 3/16 for WT), and 10/13 (77%) of TATV target sites had >20% indel for RVR (vs. 0/13 for WT). Moreover, the RR variant also had significant activity at VYCV target sites (>20% indel for 25/37 (68%) of sites) (Fig. 2h), although this activity was somewhat lower than at TYCV sites, consistent with the *in vitro* cleavage assay. We also tested the RR variant in murine Neuro2a cells (**Supplementary Fig. 1d**) and observed high rates of editing (>20% indel for 7/9 TYCV or CCCC sites).

We evaluated the genome-wide editing specificity of the RR and RVR variants using BLISS (double-strand breaks labeling *in situ* and sequencing), which quantifies DNA double-stranded breaks (DSBs) across the genome^16^. To compare the variants to WT, we restricted our analysis to target sites bearing PAMs that can be reliably cleaved by all three enzymes; TTTV was the only PAM that met this criterion, although it has lower activity for the RR variant. For three of the four target sites evaluated (*VEGFA*, *GRIN2B*, and *DNMT1*), no off-target activity was detected from deep sequencing of any of the BLISS-identified loci (Fig. 3a), either for WT or for the variants. For the fourth target site (*EMX1*), BLISS identified 6 off-target sites with detectable indels; all 6 sites had a TTCA PAM and no more than one mismatch in the first 19 bp of the guide. As expected, both variants had increased activity at these off-target sites compared to WT, consistent with their increased ability to recognize TTCA PAMs. On the other hand, when targeting a different site with known TTTV off-target sites^5^, the variants exhibited reduced off-target activity (Fig. 3b), which is also consistent with PAM preference. Collectively, these results indicate that the variants retain a high level of editing specificity that is comparable to WT AsCpf1.

**Figure 3.**
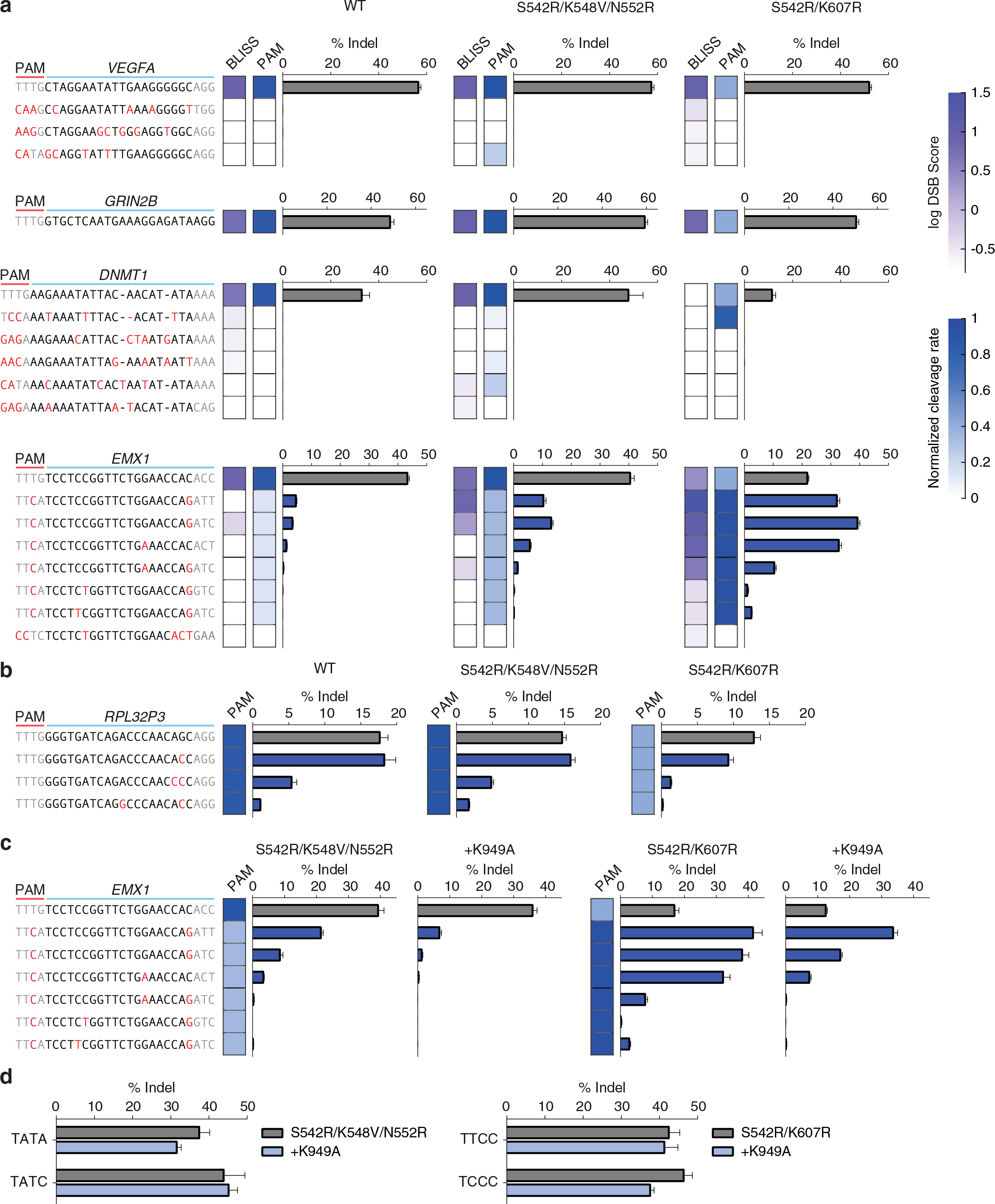
DNA targeting specificity of Cpf1 PAM variants. (**a**) DNA double-strand Breaks Labeling *In Situ* and Sequencing (BLISS) for 4 target sites (*VEGFA*, *GRIN2B*, *EMX1*, and *DNMT1*) in HEK293 cells. The log10 double-strand break (DSB) scores for BLISS are indicated by the purple heat map, and the relatively PAM cleavage rates from the *in vitro* cleavage assay are indicated by the blue heat map. Mismatches in the last three bases of the guide are not highlighted as they do not impact cleavage efficiency^4, 5^. (**b**) Evaluation of an additional target site with known TTTV off-target sites^5^, demonstrating the contribution of PAM preference to off-target activity. As the variants, particularly RR, have lower preference for TTTV PAMs compared to WT, cleavage at the TTTV off-target sites assessed was also reduced. (**c**) Addition of a K949A mutation reduces off-target DNA cleavage. (**d**) Combining K949A with the S542R/K548V/N552R and S542R/K607R PAM variants retains high levels of on-target activity for their preferred PAMs.

Finally, we investigated whether specificity can be improved by removing non-specific contacts between positively-charged or polar residues and the target DNA, similar to strategies previously employed with *Streptococcus pyogenes* Cas9 (SpCas9)^17, 18^. We identified K949A, which is located in the cleft of the protein that is hypothesized to interact with the non-target DNA strand, as a candidate (**Supplementary Fig. 1e**). When combined with the RR and RVR variants, K949A reduced cleavage at all off-target sites assessed (Fig. 3c) while maintaining high levels of on-target activity (Fig. 3d).

All mutations we identified are sufficiently close to the PAM duplex to introduce new or altered interactions with the DNA, consistent with previous work to change the PAM recognition of Cas9^6, 7^. Based on the crystal structure, we hypothesize that S542R, which is present in both the RR and RVR variants, may function by creating new electrostatic or hydrogen bonding interactions with DNA bases through its positively-charged arginine side chain (**Supplementary Fig. 1f**). These interactions may resemble the T1337R mutation in SpCas9, which creates two new hydrogen bonds with a 4^th^ guanine base in the PAM^8, 9^. It is unclear how the other mutations contribute to alteration of PAM preference in the Cpf1 variants. Further structural studies will elucidate their altered PAM recognition mechanisms.

Because Cpf1-family enzymes have strong sequence and structural homology, the S542, K548, N552, and K607 positions in AsCpf1 have unambiguous correspondences in other Cpf1 orthologs. This raises an intriguing question of whether the mutations we identified can be applied to alter the PAM preferences of other orthologs. For instance, we hypothesize that, based on the crystal structure^19^, LbCpf1 can be engineered to recognize TYCV/CCCC and TATV PAMs by the mutations G532R/K595R and G532R/K538V/Y542R, respectively.

In summary, we have demonstrated that despite strong evolutionary conservation, the PAM preference of Cpf1 can be altered by suitable mutations to residues close to the PAM duplex. The two variants of AsCpf1 we engineered can robustly cleave target sites with TYCV/CCCC and TATV PAMs, respectively, in mammalian cells. Collectively, they extend the targeting range of Cpf1 to one cleavage site for every 8.7 bp in non-repetitive regions of the human genome, which compares favorably to SpCas9. We anticipate that these variants will be a useful addition to the CRISPR-Cas genome engineering toolbox.

## Methods

*Library construction*. Human codon-optimized AsCpf1 driven by a T7 promoter was cloned into a modified pACYC backbone, and unique restriction sites were introduced flanking the selected PAM-proximal AsCpf1 residues via suitable silent mutations. For each residue, a mutagenic insert was synthesized as short complementary oligonucleotides (Integrated DNA Technologies), with the mutated codon replaced by a degenerate NNK mixture of bases (where K = G or T). Each degenerate codon position was also barcoded by creating a unique combination of silent mutations in non-mutated neighboring codons in order to correct for sequencing errors during screen readout. The variant library was assembled by cassette mutagenesis, mini-prepped, pooled, and precipitated with isopropanol.

*E. coli negative selection screen*. NovaBlue(DE3) *E. coli* expressing the T7 polymerase (Novagen) were transformed with the variant library and plated on LB agar containing 100 μg/mL ampicillin. Surviving colonies were scraped and cultured in ZymoBroth with 100 μg/mL ampicillin to an O.D. of 0.4-0.6 and made competent using a Mix & Go kit (Zymo). For each mutant PAM screened, the competent *E. coli* pool was transformed with 100 ng target plasmid containing the mutant PAM, incubated on ice for 15-30 min, heat shocked at 42 °C for 30s, and plated on LB agar (Affymetrix) containing 100 μg/mL ampicillin and 25 μg/mL chloramphenicol in the absence of IPTG. A negative control was obtained by transforming the *E. coli* with pUC19, which lacks the target site. Plasmid DNA from surviving colonies was isolated by midi-prep (Qiagen). The regions containing mutations were amplified with custom primers containing Illumina adaptors and sequenced with a paired-end 600-cycle MiSeq kit (Illumina). Reads were filtered by requiring perfect matches to silent codon barcodes; a Phred quality (Q score) of at least 30 for each of the three NNK bases; and consistency between forward and reverse reads, when applicable. The read count for each variant was normalized assuming that the mean abundance of TAG (stop) codons was equivalent to the negative control.

*In vitro PAM identification assay*. Cpf1 variants were transfected into HEK293 cells as described below. Cell lysate was prepared with lysis buffer (20 mM HEPES, 100 mM KCl, 5 mM MgCl_2_, 1 mM DTT, 5% glycerol, 0.1% Triton X-100) supplemented with ETDA-free cOmplete Protease Inhibitor Cocktail (Roche). T7-driven crRNA was transcribed *in vitro* using custom oligonucleotides and HiScribe T7 *in vitro* Transcription Kit (NEB) following the manufacturer’s recommended protocol. For the PAM library, a degenerate 8 bp sequence preceding a 33 bp target site1 was cloned into the MCS in pUC19, and the library was digested with AatII and LguI and gel extracted prior to use. Each *in vitro* cleavage reaction consisted of 1 μL 10x CutSmart buffer (NEB), 25 ng PAM library, 250 ng *in vitro* transcribed crRNA, 0.5 μL of cell lysate, and water for a total volume of 10 μL. Reactions were incubated at 37 °C and quenched by adding 50 μL Buffer PB (Qiagen) followed by column purification. Purified DNA was amplified with two rounds of PCR over 29 total cycles using custom primers containing Illumina adaptors and sequenced with a 75-cycle NextSeq kit (Illumina). For each Cpf1 variant, separate *in vitro* cleavage reactions were carried out for 1.15 min, 4 min, 10 min, 15 min, 20 min, 30 min, 40 min, 60 min, 90 min, and 175 min. A negative control, using lysate from unmodified HEK293 cells, was taken at 10 min.

*Computational analysis of PAM cleavage kinetics*. Sequencing reads were filtered by Phred quality (≥30 for all of the 8 degenerate PAM bases). For every cleavage reaction, a normalization factor α was computed by centering the read count histogram for NNNNVRRT sequences, which were not cleaved by WT AsCpf1 or any of the variants, at 1. The read counts for all 4^8^ PAM sequences in the reaction were then divided by α. For each cleavage reaction, a depletion ratio for each of the 4^8^ PAM sequences was calculated as (normalized read count in cleavage reaction) / (normalization read count in negative control). The depletion ratios of each PAM sequence (4^8^ total) across time points for each Cpf1 variant were fit using non-linear least squares to an exponential decay model × *x*(*t*) = *c*_0_ + *ce-^kt^*, where × (t) is the depletion ratio at time t, and the terms *c*_0_ ≤ 0.2, *c*, and *k* (the rate constant in min^-1^) are parameters. For each variant, the cleavage rate *k* of each 4-base PAM was computed as the median cleavage rate of the 256 8-base sequences corresponding to that PAM; for instance, the cleavage rate of TTTA was computed as the median cleavage rate of the 256 sequences of the form NNNNTTTA. Finally, all cleavage rates were adjusted such that the highest rate of any 4-base PAM was equal to 1 for each variant.

*Cell culture and transfection*. Human embryonic kidney 293 and Neuro2a cell lines were maintained in Dulbecco’s modified Eagle’s medium supplemented with 10% FBS (Gibco) at 37°C with 5% CO_2_ incubation. Cells were seeded one day prior to transfection in 24- or 96-well plates (Corning) at a density of approximately 1.2 × 10^5^ cells per 24-well or 2.4 × 10^4^ cells per 96-well and transfected at 50-80% confluency using Lipofectamine 2000 (Life Technologies), according to the manufacturer’s recommended protocol. For cell lysates, 500 ng of Cpf1 plasmid was delivered per 24-well. For indel analysis in HEK293 cells, a total of 400ng of Cpf1 plasmid plus 100ng crRNA plasmid was delivered per 24-well, or 100ng Cas9 plus 50ng crRNA plasmid per 96-well. For BLISS and for indel analysis in Neuro2a cells, 500 ng of a plasmid with both Cpf1 and crRNA were delivered per 24-well. All indel and BLISS experiments used a guide length of 23 nucleotides.

*Indel quantification*. All indel rates were quantified by targeted deep sequencing (Illumina). For indel library preparation, cells were harvested approximately 3 days after transfection, and genomic DNA was extracted using a QuickExtract DNA extraction kit (Epicentre) by re- suspending pelleted cells in QuickExtract (80μL per 24-well, or 20μL per 96-well), followed by incubation at 65°C for 15min, 68°C for 15min and 98°C for 10min. PCR amplicons for deep sequencing were generated using two rounds of PCR as previously described20. Indels were counted computationally by searching each amplicon for exact matches with strings delineating the ends of a 70bp window around the cut site. The distance in bp between these strings was then compared to the corresponding distance in the reference genome, and the amplicon was counted as an indel if the two distances differed. For each sample, the indel rate was determined as (number of reads with an indel) / (number of total reads). Samples with fewer than 1000 total reads were not included in subsequent analyses. Where negative control data is not shown, indel percentages from targeted deep sequencing represent background-subtracted maximum likelihood estimates. In particular, for a sample with *R* total reads, of which *n* ≤ *R* are indels, and false positive rate 0 ≤ α ≤ 1 (as determined by the negative control), the true indel rate was estimated as max {0,[(*n*/*R*) − *α*]/(1 − *α*).

*Computational analysis of Cpf1 targeting range*. The complete GRCh38 human genome assembly, with repeats and low complexity regions masked, was downloaded from Ensembl (ftp://ftp.ensembl.org/pub/release-86/fasta/homo_sapiens/dna/). These data indicated a total of 1.34 × 10^9^ phosphodiester bonds present in the top strand of the genome that are cleavable by at least one non-masked 23 bp Cpf1 guide (out of a possible two), regardless of the PAM. Of these bonds, 5.95 × 10^7^ are cleavable by guides with TTTV PAMs (22.6:1 ratio), while 1.55 × 108 are cleavable by guides with TTTV, TATV, TYCV, or CCCC PAMs (8.7:1 ratio).

*BLISS*. All BLISS experiments and analysis were performed as previously described^16^.

*Sample size and statistics*. The sample sizes for each measurement were *n* = 3 for bacterial colony counts (Fig. 1c); *n* = 4 combinatorial mutagenesis (Fig. 2b) and for indel analysis of BLISS loci (Fig. 3a); and *n* = 2 or *n* = 3 for all other indel data. The error bars in all figures show standard error of the mean.

## Acknowledgements

We thank A. Magnell for experimental assistance; R. Macrae for a critical review of the manuscript; and the entire Zhang laboratory for support and advice. W.X.Y. is supported by T32GM007753 from the National Institute of General Medical Sciences and a Paul and Daisy Soros Fellowship. H.N. is supported by JST, PRESTO, JSPS KAKENHI Grant Numbers 26291010 and 15H01463. O.N. is supported by the Basic Science and Platform Technology Program for Innovative Biological Medicine from the Japan Agency for Medical Research and Development, AMED, and the Council for Science, and Platform for Drug Discovery, Informatics, and Structural Life Science from the Ministry of Education, Culture, Sports, Science and Technology. F.Z. is supported by the National Institutes of Health through NIMH (5DP1- MH100706 and 1R01MH110049) and NIDDK (5R01DK097768-03), a Waterman Award from the National Science Foundation, the Keck, New York Stem Cell, Damon Runyon, Searle Scholars, Merkin, and Vallee Foundations, and B. Metcalfe. F.Z. is a New York Stem Cell Foundation Robertson Investigator. Plasmid DNA encoding AsCpf1 variants S542R/K548V/N552R and S542R/K607R are available from Addgene under a Universal Biological Material Transfer Agreement with the Broad Institute and MIT. F.Z. is a founder and scientific advisor for Editas Medicine and a scientific advisor for Horizon Discovery. Further information about the protocols, plasmids, and reagents can be found at the Zhang laboratory website (www.genome-engineering.org).

## Author Contributions

L.G., D.C., and F.Z. conceived this study. L.G. and D.C. performed experiments with help from all authors. L.G. wrote code for data analysis. J.M. contributed to the bacterial selection screen. M.S. processed BLISS samples, and W.X.Y. analyzed BLISS data. T.Y., H.N., and O.N. provided unpublished AsCpf1 crystal structure information. F.Z. supervised research. L.G. and F.Z. wrote the manuscript with input from all authors.

